# Deep-Learning Sleep Staging of Whole-Night EEG–fMRI Data Achieves Human-Level Agreement and Aligns With MRI-Defined Brain States

**DOI:** 10.64898/2026.11.24.768884

**Authors:** Fan Nils Yang, Dante Picchioni, Jeff H. Duyn

## Abstract

Concurrent EEG–fMRI recordings are increasingly used to study the neural mechanisms and benefits of sleep in humans. However, sleep staging from EEG–fMRI data remains challenging and time-consuming because MRI gradient artifacts and increased cardio-ballistic artifacts can substantially degrade EEG signal quality. Here, we used Sleepyland, a publicly available deep-learning-based tool, to demonstrate that an ensemble of four sleep-staging algorithms can achieve performance comparable to human raters without model fine-tuning. We further found that the Sleepyland sleep scores showed stronger alignment with previously identified MRI-defined sleep brain states, particularly during REM sleep, which was verified using eye-camera recordings. Together, these findings demonstrate the feasibility of using deep-learning methods to provide an automatic and possibly more objective measure of human sleep states.

## Introduction

Polysomnography (PSG), including electroencephalography (EEG), is the gold standard for human scoring sleep stages (Berry et al., 2020). However, human scoring is both time-consuming and subjective. Over the past decade, machine-learning and deep-learning algorithms have increasingly been used to address these limitations, and current top-performing algorithms can now match or exceed average human raters in both accuracy and time-efficiency (Olesen et al., 2021; Perslev et al., 2021; Phan et al., 2022; Rossi et al., 2025; Vallat & Walker, 2021). With growing interest in characterizing sleep using fMRI (Moehlman et al., 2019; Song et al., 2022; Stevner et al., 2019; Yang et al., 2024, 2025; Zou et al., 2022), it is timely to test whether algorithms developed and trained outside the scanner can be applied to the generally reduced-quality EEG data obtained in the MRI environment without further fine-tuning.

To date, most EEG–fMRI sleep studies have focused only on characterizing NREM sleep stages (Song et al., 2022; Stevner et al., 2019; Zou et al., 2022), because participants rarely reach REM sleep during typical MRI scan durations of only 1–2 hours. Nevertheless, recent work has demonstrated the feasibility of performing whole-night EEG–fMRI recordings that include multiple sleep cycles and REM sleep (Moehlman et al., 2019). This provides a valuable opportunity to evaluate how state-of-the-art deep-learning algorithms, for example those implemented through Sleepyland (Rossi et al., 2025), perform on noisy EEG data across the full complement of naturally occurring sleep cycles and to provide recommendations for future research. In addition, we propose that deep-learning algorithms may detect sleep-related features that are difficult for human raters to identify, and that these features should be consistent across modalities, i.e., EEG and fMRI. Therefore, we hypothesize that Sleepyland predicted sleep stages will show stronger alignment with hidden Markov sleep states derived from fMRI data alone (Yang et al., 2024).

## Methods

All data used in this study were collected under a human subjects research protocol approved by the National Institutes of Health Combined Neuroscience Institutional Review Board (USA; Protocol Number 16-N-0031). Written informed consent was obtained from all participants. Data acquisition was conducted as part of a previously described sleep experiment (Moehlman et al., 2019), which included two consecutive nights of concurrent EEG–fMRI recordings while participants slept inside a 3T Siemens Skyra MRI scanner for 8 hours each night. To promote a consistent sleep schedule, participants were instructed to maintain regular sleep patterns for two weeks before the experiment, and compliance was monitored using wearable devices. No sleep-deprivation procedures were used. Although eye cameras were included in the experiment, only a limited number of runs (13 out of 183) had sufficient data quality for deep-learning-based eye tracking (Syeda et al., 2024) that was used to derive eye movement strengths in the current study.

The EEG/PSG system, including the BrainCap MR electrode cap, was manufactured by Brain Products (Gilching, Germany); see Moehlman et al. (2019) for details. To ensure accurate synchronization, EEG data acquisition was timed using the 10 MHz clock from the MRI system. The Biopac system also recorded MRI volume triggers to support synchronization with peripheral physiological recordings.

Of the 16 individuals who attempted the study, 12 (8 female; aged 18–35 years) completed both nights of scanning. Each overnight session lasted approximately from 23:00 to 07:00. During each night, fMRI acquisition was intermittently interrupted by either acoustically induced or spontaneous awakenings. As a result, each night consisted of multiple experimental runs, with durations ranging from 5 minutes to 3 hours. The EEG data were divided into multiple runs corresponding to the MRI runs. Because EEG recording began slightly before the start of MRI acquisition, the manual sleep-stage scores were not aligned with the MRI TRs. Detailed preprocessing procedures for fMRI, EEG, and peripheral physiological measures have been described elsewhere (Moehlman et al., 2019). Briefly, EEG signals were corrected for MRI gradient and cardioballistic artifacts and then downsampled to 250 Hz using Analyzer software (Brain Vision, Morrisville, USA). Manual scoring was performed in 30-second epochs using central electrodes C3 and C4, alongside EOG and EMG, following established criteria with standard filters and channel references (Berry et al., 2020). The Manual scored sleep stages include wake, N1, N2, N3, REM, and undefined stage (stages that cannot be assigned due to artifacts or other reasons). ICA-based cleaning was applied to the EEG signals (Betta et al., 2021; Mensen et al., 2016; Riedner et al., 2007). In Figure S2, we show the EEG signal before and after an MRI scan start without processing, and EEG signal after processing. It is worth noting that the quality of the EOG signals remained suboptimal even after processing, and the human sleep scorers were aware of this issue when performing sleep staging.

Among the 24 nights of data collected from 12 participants across two nights each, one night (the 2nd night of one subject) was independently evaluated by two sleep scorers. The inter-rater agreement for this night was 74.8% (Moehlman et al., 2019), slightly lower than the 80–90% typically observed outside the MRI scanner (Lee et al., 2022). The two raters then jointly rescored the night to resolve any disagreements and establish a consensus (sleep-stage) score. We first applied Sleepyland to this data to obtain ensemble sleep-stage predictions (Sleepyland scores) and confidence estimates from four different algorithms (Four algorithms: YASA, U-Sleep, DeepResNet, and SleepTransformer), which have demonstrated surpasses the best human scorer (macro-F1, 85.2% vs 80.8%) (Rossi et al., 2025). Machine scoring from two distinct channel sets was compared: the central channels C3, C4, and Cz alone, and the same central channels combined with EOG channels. EEG data acquired before the onset of fMRI acquisition were discarded, thereby aligning the 30-s sleep epochs with the 3-s fMRI repetition times (TRs).

Previously, we applied a hidden Markov model (HMM) to the fMRI data (Yang et al., 2024). The HMM inference aimed to identify recurring discrete brain states, each characterized by a distinct statistical pattern. We used a Gaussian HMM implemented in the MATLAB toolbox HMM-MAR v1.0 (Vidaurre et al., 2017, 2018), where each state was modeled as a multivariate normal distribution capturing both first-order statistics, reflecting mean activity, and second-order statistics, reflecting covariance structure. State parameters were estimated at the group level, whereas state time courses were defined separately for each participant. Modules of multiple states were identified based on inter-state transition probabilities alone without inference from EEG or manual scores. These modules, as well as the individual state time courses, were partly consistent with manual scores but also provided additional information beyond conventional sleep staging, including subdivisions within N2 sleep. See Yang et. al. (2024) for details. In the present study, we compared the temporal correspondence among HMM state time courses, manual scores, and Sleepyland scores.

## Results

### Sleepyland score performance relative to consensus scoring

We first evaluated the performance of the Sleepyland scores on the night that had been independently scored by two human raters and adjudicated into a consensus sleep-stage score. Performance depended on the channel set used for prediction. The Sleepyland scores showed higher agreement with the consensus score when only central EEG channels were used than when EOG channels were included (0.757 vs. 0.574, excluding undefined sleep stage). Because the central-channel model produced the better agreement, subsequent benchmark analyses focused on predictions generated from just the EEG channels: C3, C4, and Cz.

The central-channel Sleepyland-predicted scores achieved an overall agreement of 0.757 with the consensus sleep-stage score, slightly higher than the inter-rater agreement between the two human scorers for the same night (0.748). Average confidence score for each sleep stage is Wake: 0.477, N1: 0.407, N2: 0.769, N3: 0.637, REM: 0.593. Agreement improved as low-confidence epochs were removed from the analysis **(Table S1)**. For example, applying a confidence threshold of 0.4 increased agreement to 0.801 and Cohen’s kappa to 0.729 while retaining 87% of epochs (N1 is the stage with the lowest confidence, followed by Wake). More stringent thresholds further increased raw agreement but retained fewer epochs and, at the highest threshold, produced unstable kappa estimates because only a small subset of data remained. However, this finding reflects selective retention and should not be interpreted as improved performance for complete-night scoring. Low-confidence epochs may instead be useful for identifying segments that require manual review.

### Stage-specific performance

The confusion matrix showed that ensemble performance varied across sleep stages (**Table S2)**. Predictions were strongest for N2 and N3, with most consensus-labeled N2 and N3 epochs assigned to the same stage by the ensemble. In contrast, N1 and REM showed lower stage-specific agreement. N1 was frequently confused with wake, N2, or REM, consistent with the known difficulty of reliably human-based identification of N1. REM epochs were most often correctly classified as REM but also showed substantial confusion with wake and N2, suggesting that REM detection remains more vulnerable in the scanner environment than N2 or N3 detection.

### Alignment with fMRI acquisition and MRI-defined brain states

After establishing that the central-channel ensemble without the EOG channels produced human-level agreement on the benchmark night, we applied Sleepyland to all 24 nights. Because EEG recording began several minutes before fMRI acquisition, EEG files were truncated so that 30-second sleep-stage epochs were aligned with the onset of fMRI acquisition and the 3-s fMRI repetition time. In addition, manual scores and sleepyland scores were up-sampled to 3-s resolution to match HMM states. This alignment allowed direct comparison among manual score by one of the human raters, Sleepyland scores, and MRI-derived HMM states.

Without confidence thresholding, the Sleepyland scores showed moderate agreement with manual scores for both nights examined in detail **(Table S3)**. Agreement was higher for Night 2 than for Night 1, the adaptation night. Night 1 contained proportionally more wake and N1 and less N3 and REM, which likely reduced overall performance because wake and N1 were the stages with weaker ensemble classification in the benchmark confusion matrix.

Qualitative comparison with HMM state time courses suggested that Sleepyland scores were often more consistent with MRI-defined sleep brain states and the available eye-movement video than the Manual scores. This effect was particularly apparent for REM-related patterns, where predicted REM epochs showed closer temporal correspondence with HMM states and were supported by eye-camera evidence in available high-quality runs. Several discrepancies appeared to reflect alignment effects **(Figure S1)**, ambiguous REM periods **(Figures 1–3)**, or epochs labeled as undefined by manual scoring but classified as N3 by the Sleepyland scores in a manner consistent with the HMM state structure **(Figure 4)**. Specifically, in **Figures 1** and **2**, both the HMM states and the Sleepyland predictions identified epochs as REM that PSG-based scoring did not, and the eye-camera recordings for these epochs showed large eye movements consistent with REM. **Figure 3** shows the opposite pattern: epochs that the HMM states and Sleepyland predictions did not classify as REM had been scored as REM by the human scorer, and these epochs showed low, baseline-like eye-movement activity.

**Figure 1.**
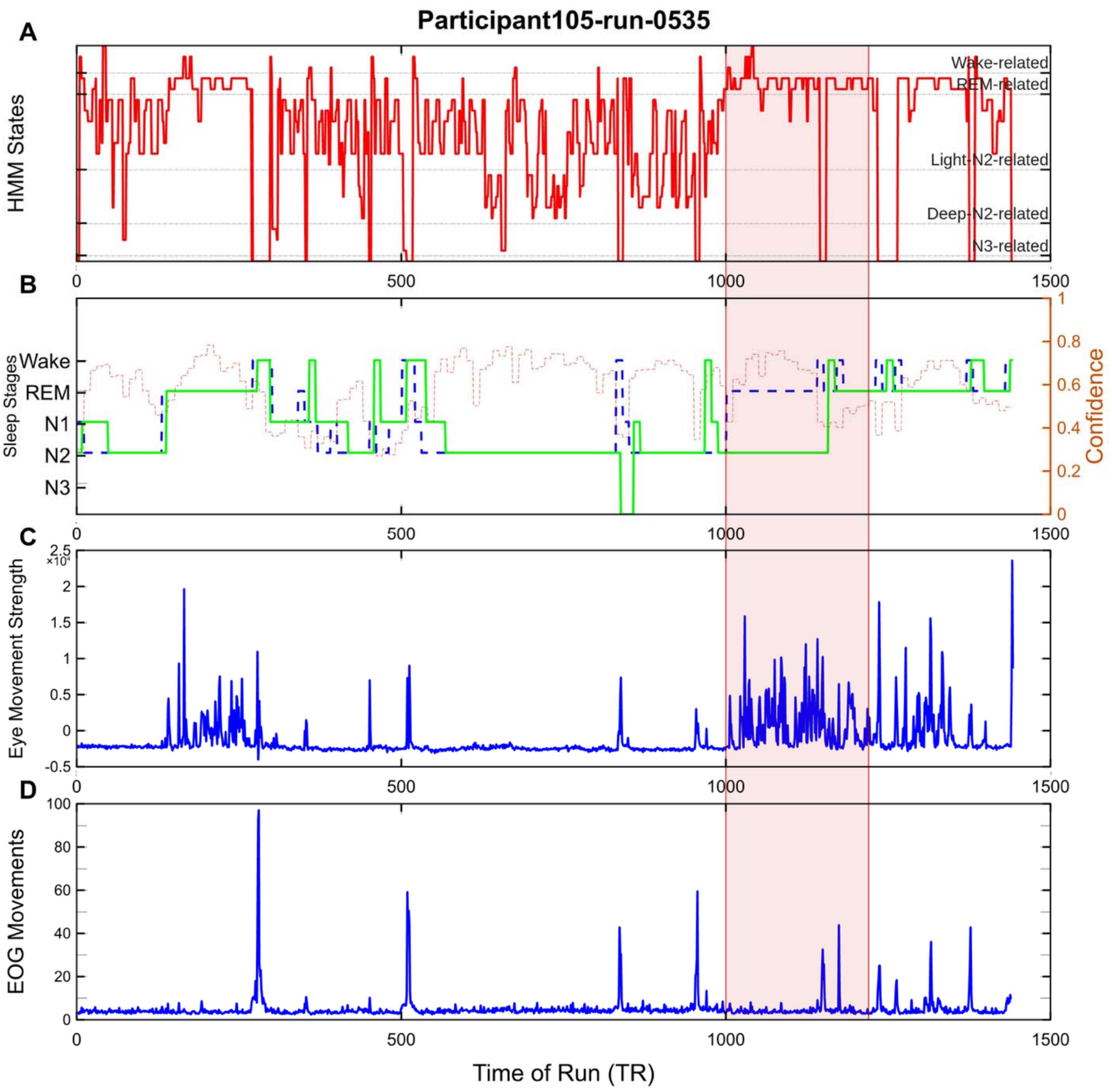
Example of REM-related discrepancy in which Sleepyland scores corresponded more closely to MRI-defined HMM states than did manual scoring. Epochs classified as REM by both the HMM and the Sleepyland scores, but not by manual scoring, coincided with large eye movements in the simultaneous eye-camera recording. Panels share the fMRI run index (Time of Run, TR) on the x-axis. In the top panel A, the red trace is the MRI-derived HMM state sequence, with states grouped along the y-axis into Wake-related, REM-related, Light-N2-related, Deep-N2-related, and N3-related clusters (state 0 means the fMRI time course is censored due to motion or an outlier). In the middle panel B, the green solid line is the manual sleep staging, the blue dashed line is the Sleepyland-predicted staging (left y-axis: Wake, REM, N1, N2, N3), and the red dashed line is the Sleepyland ensemble confidence (right y-axis, 0–1). In the panel C, the blue trace is eye-movement strength derived from the eye-camera recording. In the panel D, the blue trace is standard deviation of EOG signal per TR, used as a measure of EOG-based eye movement.

**Figure 2.**
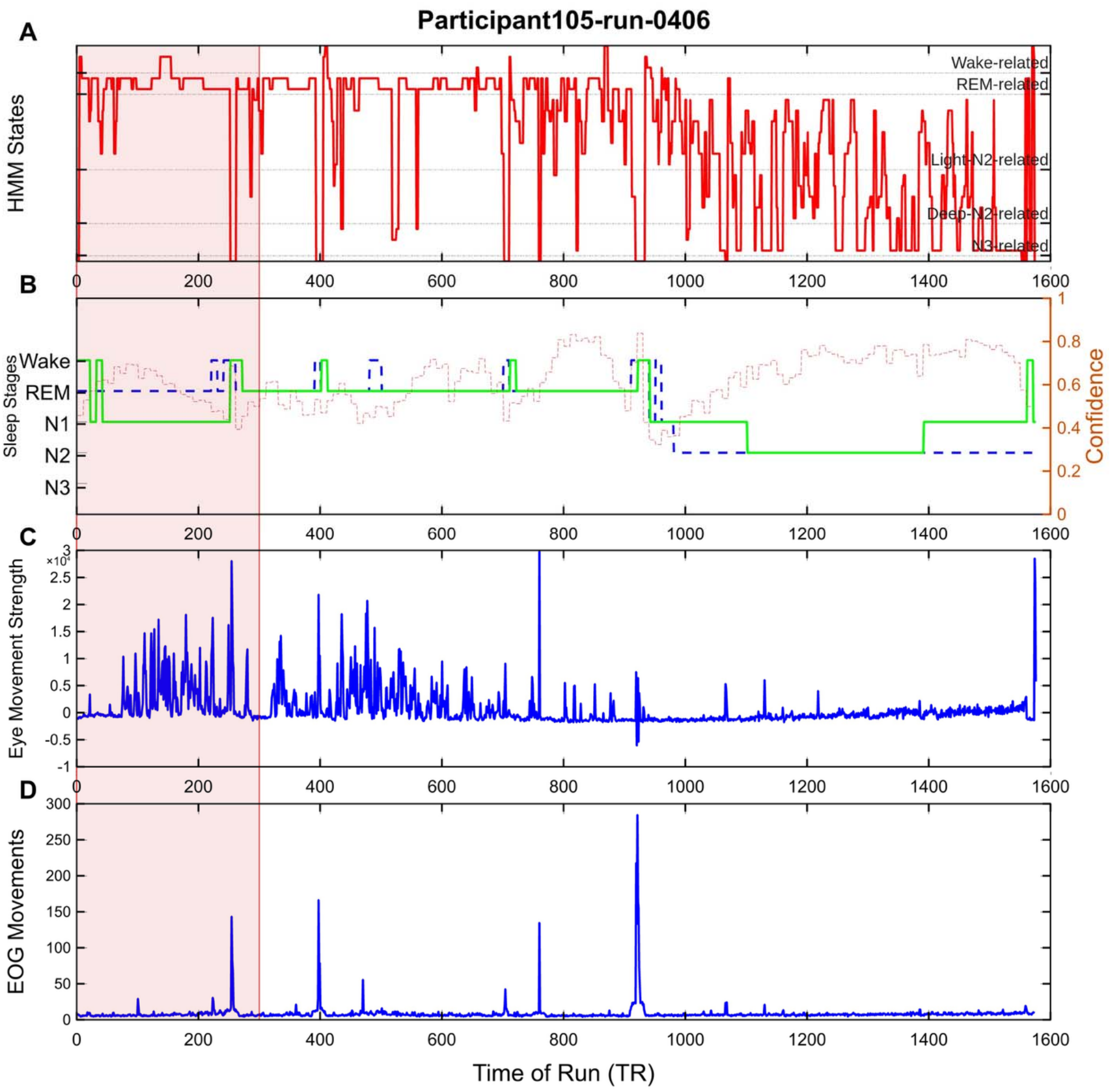
An additional example in which epochs labeled REM by the HMM states and the Sleepyland ensemble, but not by manual scoring, aligned with periods of large eye movements consistent with REM sleep. Panels share the fMRI run index (Time of Run, TR) on the x-axis. In the top panel A, the red trace is the MRI-derived HMM state sequence, with states grouped along the y-axis into Wake-related, REM-related, Light-N2-related, Deep-N2-related, and N3-related clusters (state 0 means the fMRI time course is censored due to motion or an outlier). In the middle panel B, the green solid line is the PSG-based manual sleep staging, the blue dashed line is the Sleepyland-predicted staging (left y-axis: Wake, REM, N1, N2, N3), and the red dashed line is the Sleepyland ensemble confidence (right y-axis, 0–1). In the bottom panel C, the blue trace is eye-movement strength derived from the eye-camera recording.

**Figure 3.**
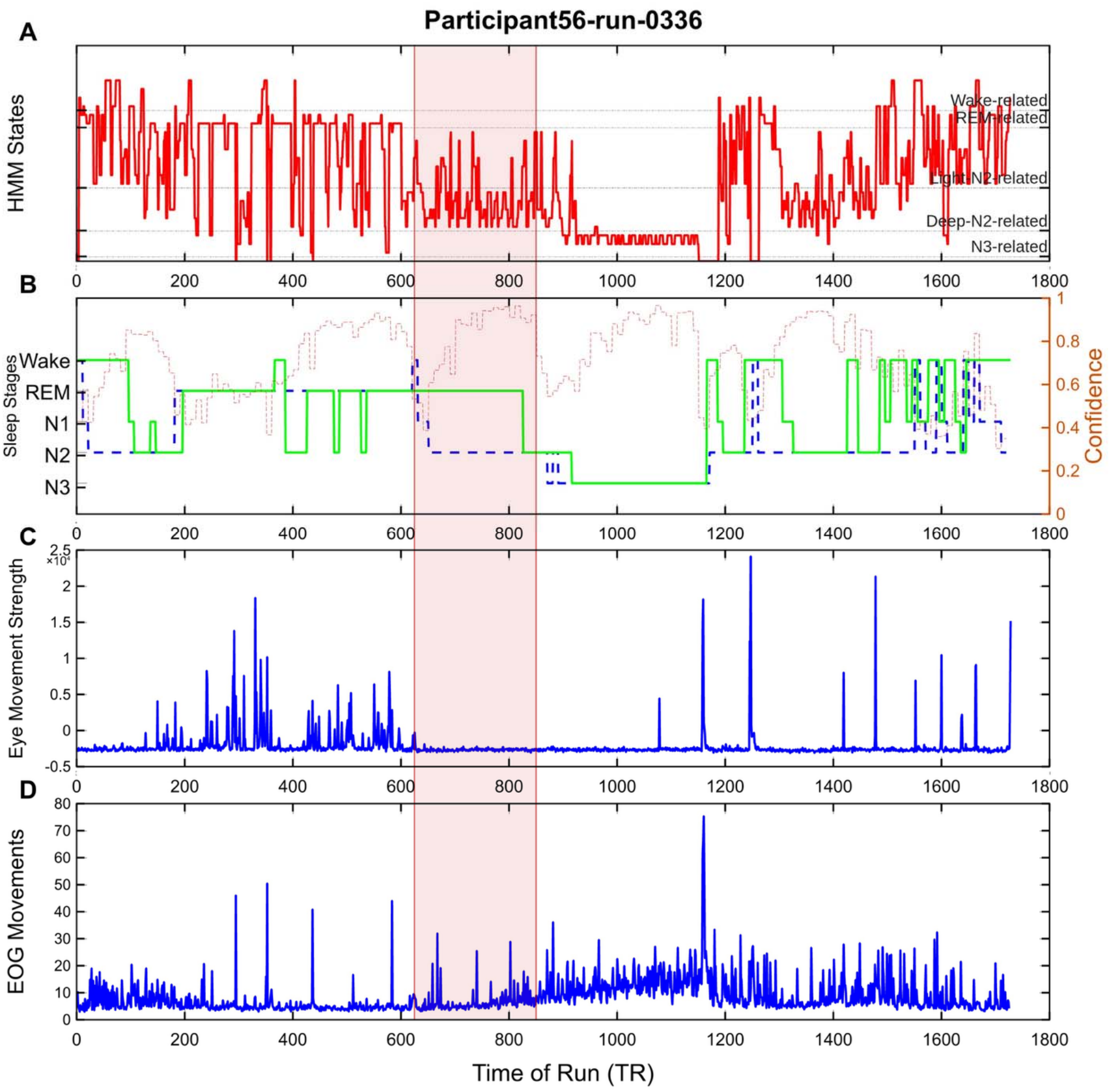
Example in which epochs scored as REM by manual scoring, but not by the HMM states or the Sleepyland ensemble, showed low, baseline-like eye-movement activity. Panels share the fMRI run index (Time of Run, TR) on the x-axis. In the top panel A, the red trace is the MRI-derived HMM state sequence, with states grouped along the y-axis into Wake-related, REM-related, Light-N2-related, Deep-N2-related, and N3-related clusters (state 0 means the fMRI time course is censored due to motion or an outlier). In the middle panel B, the green solid line is the manual sleep staging, the blue dashed line is the Sleepyland-predicted staging (left y-axis: Wake, REM, N1, N2, N3), and the red dashed line is the Sleepyland ensemble confidence (right y-axis, 0–1). In the panel C, the blue trace is eye-movement strength from the eye-camera recording. In the panel D, the blue trace is standard deviation of EOG signal per TR, used as a measure of EOG-based eye movement.

**Figure 4.**
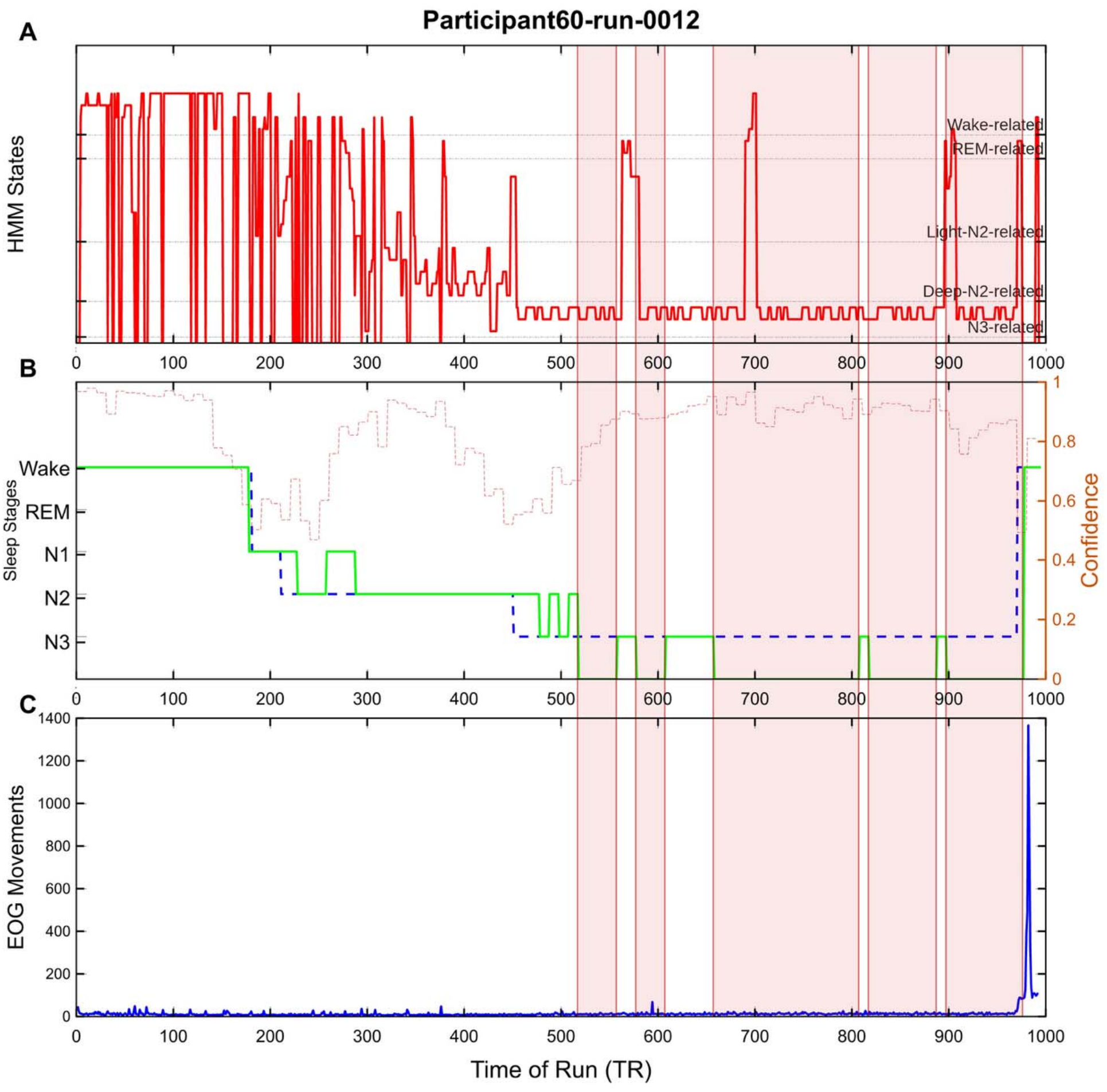
Epochs scored as undefined by manual scoring (0 on the Y axis in Panel B) but classified as N3 by the Sleepyland ensemble, consistent with the corresponding MRI-defined HMM state. Both panels share the fMRI run index (Time of Run, TR) on the x-axis. In the top panel A, the red trace is the MRI-derived HMM state sequence, with states grouped along the y-axis into Wake-related, REM-related, Light-N2-related, Deep-N2-related, and N3-related clusters (state 0 means the fMRI time course is censored due to motion or an outlier). In the panel B, the green solid line is the manual sleep staging and the blue dashed line is the Sleepyland-predicted staging (left y-axis: Wake, REM, N1, N2, N3), and the red dashed line is the Sleepyland ensemble confidence (right y-axis, 0–1); epochs left undefined by PSG (0 on the Y axis) fall below the N3 level. In the panel C, the blue trace is standard deviation of EOG signal per TR, used as a measure of EOG-based eye movement.

## Discussion

In this study, we tested whether deep-learning sleep-staging algorithms developed outside the scanner can be applied to whole-night EEG–fMRI recordings without additional fine-tuning. Using Sleepyland, an ensemble of four algorithms, we found that automated staging of artifact-corrected in-scanner EEG reached good agreement with a human consensus score. Moreover, the Sleepyland scores aligned more closely than the original manual labels with independently derived, fMRI-defined brain states, an effect that was especially clear for REM sleep and was corroborated by eye-camera recordings. Together, these results indicate that off-the-shelf deep-learning models can provide fast, objective, and physiologically meaningful sleep staging in the challenging EEG–fMRI setting.

EEG acquired during fMRI is substantially noisier than EEG recorded outside the scanner, primarily because of gradient and cardioballistic artifacts that persist even after correction. This degradation makes manual sleep scoring slow and error-prone, as reflected in the relatively low inter-rater agreement of 74.8% for our benchmark night, compared with the 80–90% typically reported outside the scanner (Lee et al., 2022). At the same time, whole-night EEG–fMRI datasets remain scarce and had relatively lower data-quality compared EEG collected without MRI interference, so training or fine-tuning a scanner-specific staging model is not currently practical. Applying algorithms pretrained on large out-of-scanner datasets therefore offers an attractive alternative, and our results show that these models generalize to the scanner environment despite the pronounced domain shift.

Sleepyland scores performed better when only central EEG channels were used than when EOG channels were added (0.757 vs. 0.574). This most likely reflects the poor quality of EOG signals in the scanner, where eye channels are especially susceptible to gradient and motion artifacts, so that including them introduced more noise than information. Notably, the algorithms still identified REM sleep from central EEG alone, without access to EOG-derived eye-movement information. This indicates that the models capture EEG signatures of REM, such as characteristic low-amplitude, desynchronized activity, rather than relying solely on the ocular features that human scorers traditionally emphasize.

Sleepyland scoring offers several practical advantages over manual PSG scoring in the EEG– fMRI context. First, it is fast and more consistent/reproducible, removing the labor and subjectivity of visual scoring. Second, it is flexible: the scoring window can be repositioned easily, for example, to align epochs with the onset of fMRI acquisition. Third, each epoch carries a confidence estimate, allowing uncertain epochs to be further human review. This is particularly useful for stages such as N1, whose inter-rater reliability is already low among human scorers (Lee et al., 2022). Fourth, the Sleepyland scores corresponded more closely than the manual scores to MRI-defined brain states, indicating that the models can track sleep-related structure that human raters may miss.

The closer correspondence between the Sleepyland scores and the MRI-defined HMM states is notable because they were derived independently from EEG and fMRI data, respectively, and therefore cannot be attributed to shared input signals. Together with the supporting eye-camera evidence, this cross-modal convergence suggests that some apparent algorithmic errors may instead reflect ambiguous or inaccurate manual scores. The clearest examples involved REM sleep: as is readily apparent in the eye-movement strength traces, epochs that both the HMM and the Sleepyland scores identified as REM but that Manual scores did not coincide with visibly elevated eye movements, whereas epochs scored as REM by Manual scores showed only baseline-level eye movement **(Figures 2–4)**. The ensemble also assigned stages to epochs that manual scoring had left undefined, in a manner consistent with the concurrent HMM state **(Figure 5)**. Sleepyland scoring may therefore help recover otherwise unusable epochs and resolve ambiguous ones, increasing the amount of analyzable data in EEG–fMRI sleep studies.

Several limitations should be acknowledged. The benchmark comparison rested on a single night scored by two raters, and the broader comparison with MRI-defined states was qualitative rather than quantitative. The dataset, although unique in providing whole-night, multi-cycle EEG–fMRI recordings, is modest in size (12 participants), which constrains generalizability. Promising next steps include domain adaptation or light fine-tuning of pretrained models on in-scanner data, improved in-scanner eye monitoring, and the use of confidence-weighted predictions as a principled way to incorporate automated sleep staging into fMRI analyses of sleep.

## Conclusion

In conclusion, off-the-shelf deep-learning Sleepyland scores can stage sleep from whole-night EEG–fMRI recordings at close to than human-level reliability without EOG channels, especially for REM stage, and can recover physiologically meaningful structure that manual scoring misses. For future EEG–fMRI sleep studies, we therefore recommend using pretrained ensemble algorithms as a fast and objective complement to, or replacement for, manual scoring; favoring central EEG channels over artifact-prone in-scanner EOG; and aligning the scoring window to the onset of fMRI acquisition. Adopting these practices should make sleep staging in the scanner faster, more reproducible, and better matched to the fMRI signal it is intended to describe.

## Acknowledgments

This research was supported by the Intramural Research Program of the National Institutes of Health (NIH). The contributions of the NIH author(s) were made as part of their official duties as NIH federal employees, are in compliance with agency policy requirements, and are considered Works of the United States Government. However, the findings and conclusions presented in this paper are those of the author(s) and do not necessarily reflect the views of the NIH or the U.S. Department of Health and Human Services. Sussan Guttman, Steve Newman are acknowledged for their advice and assistance. Xufu Liu and Xiao Liu are acknowledged for their assistance with eye video preprocessing.

## Author Contributions

F.N.Y. conceptualized the study, analyzed the data, generated figures, and wrote the original draft. D.P. collected the data and edited the manuscript. J.H.D. supervised the study and edited the manuscript.

## Financial Disclosure

The authors declare no competing financial interests.

## Non-financial Disclosure

The authors declare no competing non-financial interests.

## Data and code availability statement

The datasets are available at https://openneuro.org/datasets/ds005127/versions/1.0.4.

## Notes

### Competing Interest Statement

The authors have declared no competing interest.

