## Supplement for "Deep-Learning Sleep Staging of Whole-Night EEG–fMRI Data Achieves Human-Level Agreement and Aligns With MRI-Defined Brain States"

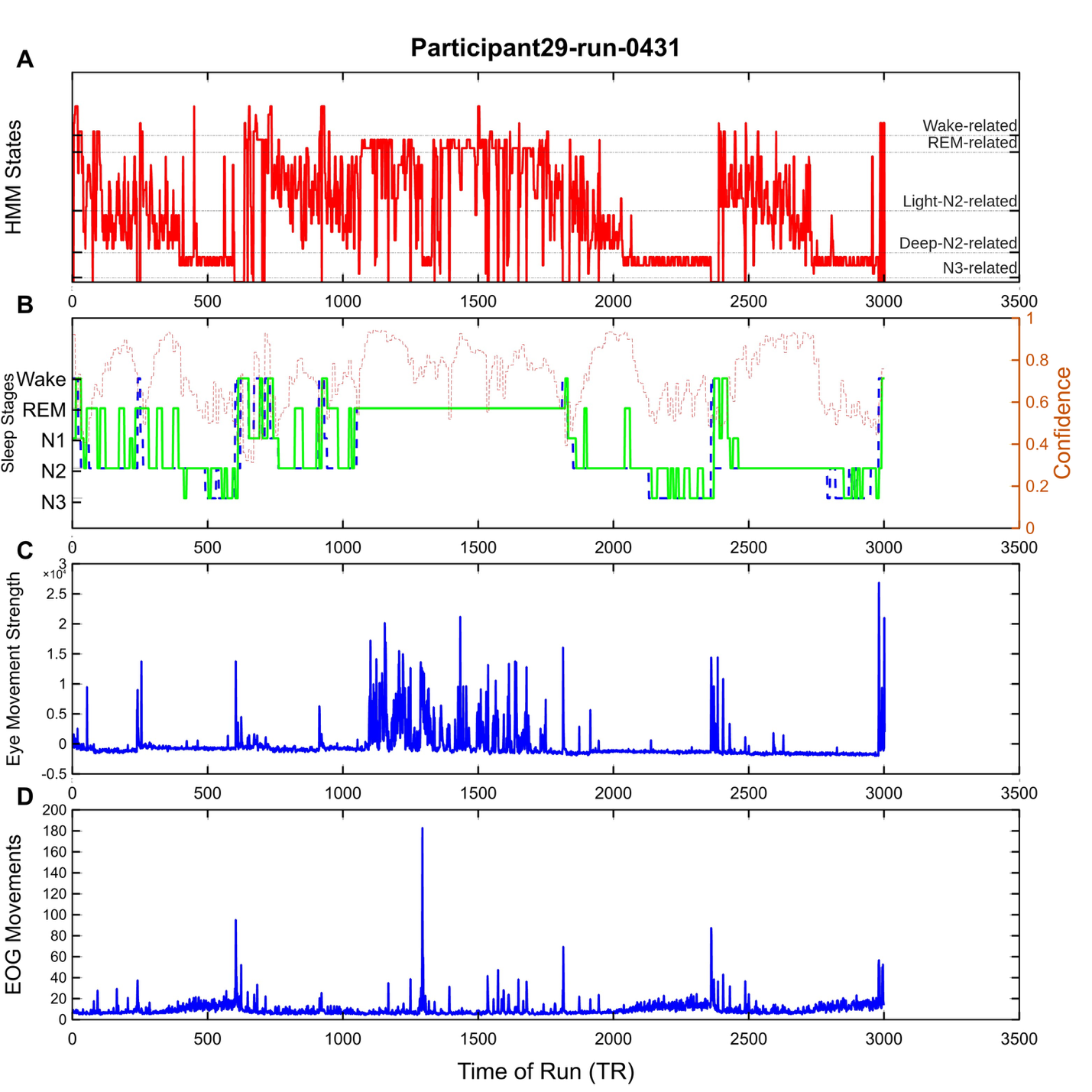

***Figure S1.*** *Effect of EEG–fMRI temporal alignment on the correspondence among manual scores, Sleepyland scores, and MRI-derived HMM state time courses. Panels share the fMRI run index (Time of Run, TR) on the x-axis. In the top panel A, the red trace is the MRI-derived HMM state sequence, with states grouped along the y-axis into Wake-related, REM-related, Light-N2-related, Deep-N2-related, and N3-related clusters (state 0 means the fMRI time course is censored due to motion or an outlier). In the middle panel B, the green solid line is manual sleep staging, the blue dashed line is the Sleepyland-predicted staging (left y-axis: Wake, REM, N1, N2, N3), and the red dashed line is the Sleepyland ensemble confidence (right y-axis, 0–1). In the panel C, the blue trace is eye-movement strength derived from the eye-camera recording. In the panel D, the blue trace is standard deviation of EOG signal per TR, used as a measure of EOG-based eye movement.*

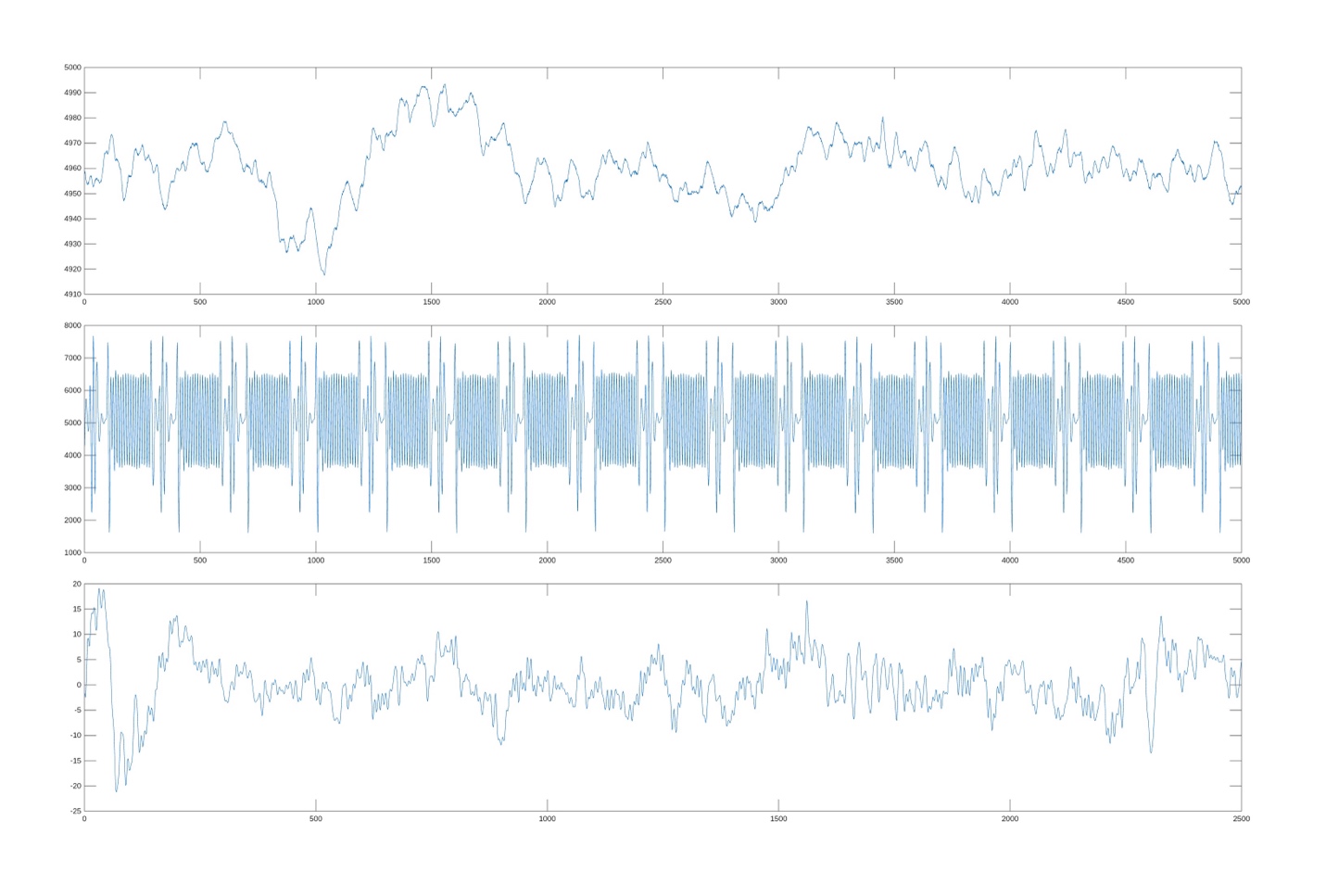

**Figure S2**. EEG data without MRI artifact (top panel), EEG data with MRI artifact (middle panel), and preprocessed EEG data (Bottom panel, removed MRI artifact, cardioballistic artifact, and with ICA cleaning).

**Table S1**. *Agreement Between Consensus Manual Scores and Sleepyland Scores Across Confidence Thresholds*

| **Confidence threshold** | **Agreement** | **Cohen's kappa** | **Data retained (%)** |
| --- | --- | --- | --- |
| 0.20 | 0.757 | 0.671 | 100 |
| 0.30 | 0.762 | 0.677 | 98 |
| 0.40 | 0.801 | 0.729 | 87 |
| 0.50 | 0.859 | 0.782 | 72 |
| 0.60 | 0.912 | 0.843 | 57 |
| 0.70 | 0.927 | 0.833 | 44 |
| 0.80 | 0.945 | 0.791 | 31 |
| 0.90 | 0.953 | 0.159 | 13 |

*Note.* Agreement and Cohen's kappa were calculated after excluding epochs with ensemble confidence below each threshold. The decrease in kappa at the highest threshold likely reflects the small and imbalanced set of retained epochs.

**Table S2**. *Confusion Matrix Comparing Consensus Manual Scores With Sleepyland Scores*

| **Consensus stage** | **Wake** | **N1** | **N2** | **N3** | **REM** |
| --- | --- | --- | --- | --- | --- |
| Wake | 87 | 6 | 17 | 0 | 8 |
| N1 | 34 | 32 | 21 | 0 | 24 |
| N2 | 6 | 2 | 298 | 1 | 4 |
| N3 | 0 | 0 | 30 | 126 | 0 |
| REM | 18 | 6 | 25 | 0 | 86 |

*Note.* Rows indicate consensus manual scores, and columns indicate Sleepyland scores

**Table S3**. *Agreement Between fMRI-Aligned Sleepyland scores and Manual Scores*

| **Night** | **Accuracy** | **Cohen's kappa** | **Confidence thresholding** |
| --- | --- | --- | --- |
| Night 2 | 0.693 (0.059) | 0.545 (0.099) | None |
| Night 1 | 0.642 (0.115) | 0.461 (0.141) | None |

*Note.* Metrics were calculated after shifting sleep-stage epochs to align with the onset of fMRI acquisition. Mean (standard deviation) of 12 participants showed here.
